# Invasion of homogeneous and polyploid populations in nutrient-limiting environments

**DOI:** 10.1101/2020.04.15.041566

**Authors:** Gregory J. Kimmel, Mark Dane, Laura Heiser, Philipp M. Altrock, Noemi Andor

## Abstract

Breast cancer progresses in a multistep process from primary tumor growth and stroma invasion to metastasis. Progression is accompanied by a switch to an invasive cell phenotype. Nutrient-limiting environments promote chemotaxis with aggressive morphologies characteristic of invasion. It is unknown how co-existing cells differ in their response to nutrient limitations and how this impacts invasion of the metapopulation as a whole. We integrate mathematical modeling with microenvironmental perturbation-data to investigate invasion in nutrient-limiting environments inhabited by one or two cancer cell subpopulations. Hereby, subpopulations are defined by their energy efficiency and chemotactic ability. We estimate the invasion-distance traveled by a homogeneous population. For heterogeneous populations, our results suggest that an imbalance between nutrient efficacy and chemotactic superiority accelerates invasion. Such imbalance will spatially segregate the two populations and only one type will dominate at the invasion front. Only if these two phenotypes are balanced do the two subpopulations compete for the same space, which decelerates invasion. We investigate ploidy as a candidate biomarker of this phenotypic heterogeneity to discern circumstances when inhibiting chemotaxis amplifies internal competition and decelerates tumor progression, from circumstances that render clinical consequences of chemotactic inhibition unfavorable.

**Significance:** A better understanding of the nature of the double-edged sword of high ploidy is a prerequisite to personalize combination-therapies with cytotoxic drugs and inhibitors of signal transduction pathways such as MTOR-Is.

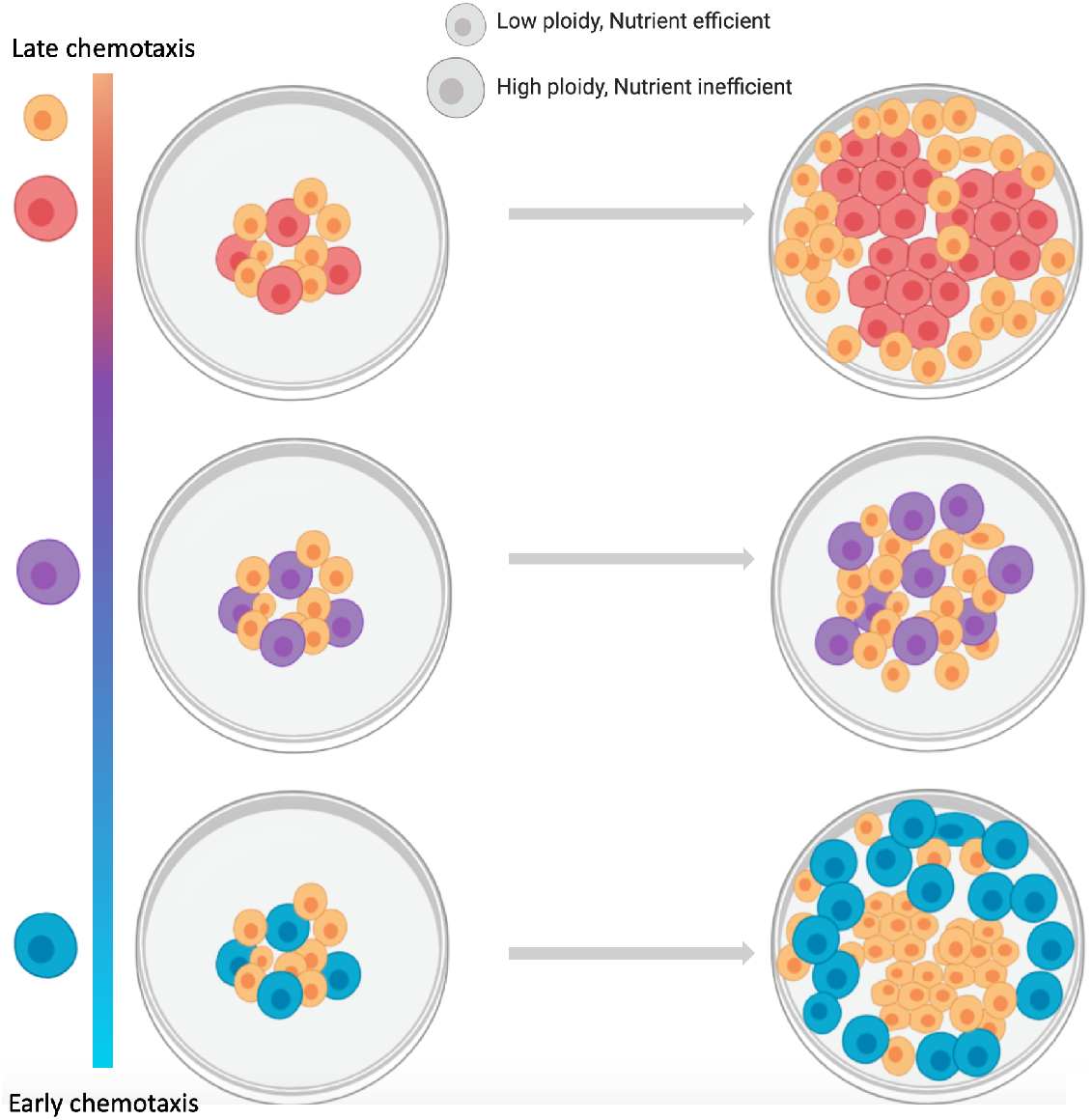

## Introduction

Invasion and infiltration are hallmarks of advanced cancers, including breast cancer, and accumulating evidence suggests that invasive subclones arise early during tumor evolution [1].

Infiltrating and invasive phenotypes are often observed among high-ploidy cells. Converging evidence from different cancer types, including colorectal-, breast-, lung- and brain cancers, suggests a strong enrichment of high ploidy cells among metastatic lesions as compared to the primary tumor [2, 3]. Even in normal development: trophoblast giant cells - the first cell type to terminally differentiate during embryogenesis - are responsible for invading the placenta and these cells often have hundreds of copies of the genome [4]. Coexistence of cancer cells at opposite extremes of the ploidy spectrum occurs frequently in cancer and is often caused by whole genome doubling (WGD). Similar to infiltration, the timing of WGD is early in tumor progression across several cancer types [5, 6], including breast cancer. Tetraploid cells resulting from WGD often lose regions of the genome, giving rise to poly-aneuploid cancer cells (PACCs). Multiple studies have described a minority population of PACCS with an unusual resilience to stress [7–9]. A very recent investigation of evolutionary selection pressures for WGD suggests that it mitigates the accumulation of deleterious somatic alterations [10]. However, it is not clear what costs cells with a duplicated genome pay for this robustness.

To address this question, we developed a mathematical model of co-evolving high- and low-ploidy clones under various energetic contingencies. We calibrate the model to recapitulate doubling times and spatial growth patterns measured for the HCC1954 ductal breast carcinoma cell line via MEMA profiling [11]. This includes exposure of HCC1954 cells to HGF in combination with 48 extracellular matrices (ECMs), followed by multi-color imaging [12]. Our results show that long-term coexistence of low- and high-ploidy clones occurs when sensitivity of the latter to energy scarcity is well-correlated to their chemotactic ability to populate new terrain. Higher energy uniformity throughout population expansion steers selection in favor of the low-ploidy clone, by minimizing the fitness gain the high-ploidy clone gets from its chemotactic superiority. Better understanding of how these two phenotypes co-evolve is necessary to develop therapeutic strategies that suppress slowly-proliferating, invasive cells before cytotoxic therapy favors them.

## Materials and Methods

We first introduce the conceptual framework that lead to the model, formulate the model equations and then we derive analytical and numerical solutions. Finally we describe drug-sensitivity and RNA sequencing data analysis of cell lines with different ploidies.

### Overall model design

The use of partial differential equations (PDEs) over a stochastic-based approach such as agent-based modeling permits us to make predictions based on analytical results derived from the subsequent PDEs and an increase in computational efficiency.

We modeled growth dynamics in polyploid populations of various subpopulation compositions. Appealing to a continuity description and assuming a continuum approximation of the cellular and energy concentration is valid, we derived a system of coupled PDEs. Each compartment in the PDE describes the spatio-temporal dynamics of the quantity of interest (e.g. energy or cellular dynamics). At the core of our model lies the assumption that chemotactic response to an energy gradient is a function of the cell’s energetic needs. This trade-off implies that heterogeneous populations will segregate spatially, with higher energy-demanding cells leading the front of tumor growth and invasion. In contrast, for an energy-rich environment we expect the cells to grow in a similar way as they will have no need to search for places of higher energy density.

We model competition for energy in a heterogeneous population, consisting of goer and grower subpopulations, to predict their behavior during plentiful and energy sparse conditions. We assume both goer and grower have the same random cell motility coefficient Γ, the same chemotactic coefficient χ and maximal growth rate λ. Sensitivity to the available energy is modeled via a Michaelis-Menten type equation with coefficients Φ_*i*_ that determine the amount of energy that population *i* needs for a half-maximal growth rate. Chemotactic motion is asymmetrically sensitive to the amount of energy available. This is accounted for by Ξ_*i*_. The goers (*U*) are more motile and require more energy compared to the growers (*V*). This manifests itself mathematically via the parameter relations Φ_*U*_ > Φ_*V*_ and Ξ_*U*_ < Ξ_*V*_.

Quantitative estimates of how a cell’s growth rate and motility depends on energy availability have been described [13–15]. Energetic resources come in various forms, and the identities of the limiting resources that ultimately drive a cell’s fate decision vary in space and time. We used MEMA profiling to investigate what likely is only a narrow range of that variability – 48 HGF-exposed ECMs [12]. HGF stimulates both growth and migration of epithelial and endothelial cells in a dose-dependent manner, whereby maximal growth-stimulating effects have been reported at concentrations twice as high as concentrations that maximize migration [16]. In line with these reports, our model demonstrates a shift from proliferation to migration as resources get depleted.

Mathematical models of a dichotomy between proliferation and migration are numerous [17–19], but whether the two phenotypes are indeed mutually exclusive remains controversial [20]. Our efforts to use mathematical modeling to inform what cost high-ploidy cells (goers) pay for their robustness builds upon these prior works. We extend it by accounting for differences in the rate at which cells consume energy (*δ*) and differences between media in the rate at which energy diffuses (coefficient Γ_*E*_). For mid-range energy diffusion coefficients our model describes directed cell motility in response to a gradient of a soluble attractant, i.e. chemotaxis. By contrast, small values of Γ_*E*_ approximate cell motility towards insoluble attractants, i.e. haptotaxis. As such, the chosen value for Γ_*E*_ sets where along the continuum between haptotaxis and chemotaxis directed cell movement resides. A special case applies when the energy diffusion coefficient is very large relative to cell movement, in which case neither chemotaxis nor haptotaxis occurs. All these energetic contingencies determine whether phenotypic differences between goers and growers manifest as such, and explain why nonproliferative arrested cells can have the same motility as cycling cells [20].

### Quick guide to equations

Our model assumes that the energy diffusion coefficient Γ_*E*_ depends on the type of media or surface upon which the cells grow. We also suppose that energy is consumed in proportion to the amount of cells present. For cell motility, we assume it is driven both by random cell motion and chemotaxis. This leads to our general coupled system:

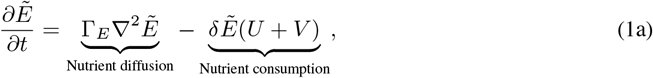

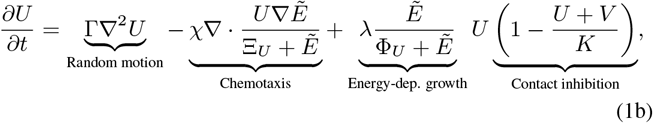

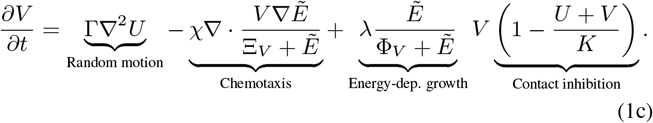

The system (1) is defined on a dish of radius *R* subject to no-flux boundary conditions. We assume the goers (*U*) and growers (*V*) are initially concentrated at the center with radius *ρ*_0_ ≤ *R* and initial concentrations *U*_0_, *V*_0_. Further, we assume that the energy density is uniformly distributed on the plate with initial value *E*_0_. All parameters except otherwise stated are independent of any of the state variables 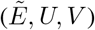. The energy is consumed at rate *δ*. Both cells can divide at the maximal rate λ, but are restricted by the energy density 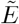. The cells can locally grow to a local maximal density given by *K*. This parameter is often cell line-dependent and is related to contact inhibition and a cell’s ability to grow on top of each other.

We now convert our system to dimensionless form that was used for all subsequent simulations and analysis via appropriate re-scaling (Supplementary methods),

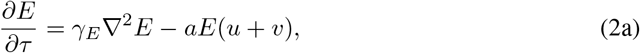

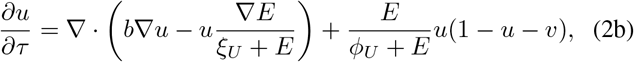

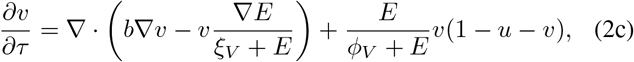

Hereby re-scaling simplified the system from thirteen (ten parameters and three initial conditions) to nine (seven parameters and two initial conditions), with dimensionless variables *γ_E_* = Γ_*E*_/*χ*, *a* = *δK*/λ, *b* = Γ/*χ*, *ϕ* = Φ/*E*_0_ and *ξ* = Ξ/*E*_0_.

### Analytical estimates of infiltration

The desire for cells to move is inherently tied to the availability of nutrients and space. To this end we define Φ(*τ*):= [*ρ*(*τ*) – *ρ*_0_]/*ρ*_0_ where *ρ*(0) = *ρ*_0_, the initial radius of cell seeding density. Φ can be thought of as a non-dimensional measure of infiltration attained after time *τ*. This dimensionless measure has the added benefit of being scale-independent. An inherent difficulty with random cell motility and calculating infiltration is that the system always reaches the boundary of the dish in finite time. Instead we will define the maximum degree of infiltration to be given by the time needed for the total energy to be below a threshold *ε* ≪ 1.

In general, the maximum degree of infiltration is difficult to predict analytically, so we only considered the single subpopulation case when obtaining our analytical estimates. We also made use of the simplification that most energy-type molecules (e.g. glucose) have a diffusion coefficient that is very large, relative to cell movement. This allows us to write a reduced model which has energy homogeneous in space (see (S3a)-(S3b) in Supplementary Methods). We estimate infiltration analytically through two different approaches. First, we extract an ODE system that couples nutrient consumption to the wave front location. Second, we derive estimates of the infiltration achieved by using the cell concentration after it has become uniform.

#### No chemotaxis infiltration estimate

We can derive an estimate for infiltration in the absence of chemotaxis by appealing to (S3a)-(S3b). These logistic growth-reaction-diffusion models often exhibit complex dynamics. One such example is that of a traveling-wave solution, where one state, typically the stable state, travels (infiltrates) through the domain. The canonical example of this phenomenon is the Fisher-KPP. In contrast to the Fisher-KPP and other traveling-wave problems typically studied, our model has a decaying growth rate and so the magnitude of the non-linearity that caused the traveling wave is tending to zero. Therefore, in the classic sense, our system does not admit a traveling wave. We here extend the theory by assuming a separation of time scales between consumption of energy (e.g. decay of energy-dependent growth rate) and the speed of the traveling wave.

To begin, we make the assumptions that the wave speed is a slow function of *r* and *τ*. The solution obtained will verify that these assumptions are valid for our system. Our ansatz takes the form *u*(*r, τ*) = *U*(*r* - *ητ*) = *U*(*z*). Note that in spatial equilibrium, *u* = 1 is stable and *u* = 0 is an unstable steady state. If the unstable state is what governs the wave speed, then the wave is said to be “pulled”, otherwise it is “pushed” [21, 22]. The resulting analysis yields a coupled system of ODEs that govern the speed of the front (Supplementary methods):

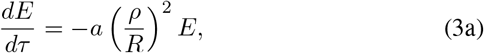

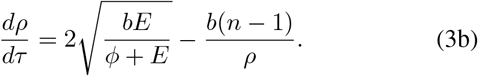

We see that our assumptions on the behavior of the traveling wave are verified, since *ρ* is assumed much larger than 0 and 0 ≪ *ρ* ≪ *R*. This shows that *E* is a slowly varying function of time and *η* = *dp*/*dτ* is a slowly varying function of time and its current distance from the center. In other words, a traveling wave will only form when the initial seeding radius is relatively large (*ρ*_0_ ~ 1 was sufficient in most simulations).

#### Estimating the degree of infiltration from equilibration

The previous section yields a system that can be integrated to track the evolution of the cell front over time. However, we may be more interested in how far it will ultimately travel before energy exhaustion and not the speed at which it gets there. An interesting alternative to tracking the wave over time is to only assume it travels as a wave, but only record the density after the system has reached uniformity. This is possible if the death rate (which has been neglected) is much smaller than the time it would take the cells to spread uniformly. If this is the case, we can bound the degree of infiltration from only knowing the uniform value u at the end of the experiment (see Supplementary Methods for details):

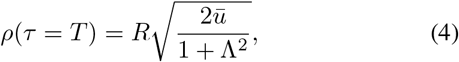

where Λ is the transition width, i.e. the length scale on which cell concentration goes from u = 1 to *u* = 0, and *ū* it is the concentration at the end of the experiment (at equilibration). Solving (4) gives us the estimated infiltration as:

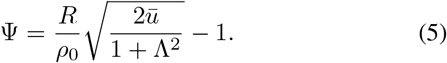

Since we assume that the transition width is unknown, we can bound Φ (or *ρ*(*T*)) by considering the lower and upper bounds Λ = 0, 1, respectively.

### Numerical estimates of infiltration

When directed cell movement is not negligible, analytical approximations are more difficult to obtain, and numerical simulations are preferred to estimate the degree of infiltration. For simulations where we measured infiltration, we took the 1-norm 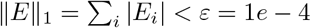 as the threshold defining minimum energy requirements. We calibrate the model to recapitulate doubling times and spatial growth patterns measured for the HCC1954 ductal breast carcinoma cell line via MEMA profiling. The dataset includes exposure of HCC1954 cells to HGF in combination with 48 ECMs in a DMSO environment (i.e. no drug was added to the media). Between 13 and 30 replicates of each ECM are printed on a rectangular MEMA array as circular spots (Supplementary Fig. 1A), adding up to a total of 692 spots [12].

#### MEMA data analysis

An average of 31 cells (90% confidence interval: 23-41) were seeded on each 350 *μm* spot and grown for three days. The median number of cells imaged at day three was 121 – which falls within the range expected from the estimated seeding density and a doubling time of approximately 43.81 hours reported for this cell line (72-128 cells). Confluence at seeding was calculated from the ratio between the cumulative area of cells and the area of the spot (see Supplementary Fig. 1B-C).

Quantification of segmented, multi-color imaging data obtained for each spot three days post-seeding was downloaded from Synapse (synID: syn9612057; plateID: LI8C00243; well: B03). Cells tended to grow in a symmetric, toroidal shape (Supplementary Fig. 1D), albeit considerable variability was observed across the ECMs. We binned cells detected within a given spot according to their distance to the center of the spot and calculated the confluence of each bin (Supplementary Methods). This was then compared to the confluence obtained from the simulations as described below.

#### Simulation environment

Simulations were ran to recapitulate logistics and initial growth conditions on the MEMA array (Supplementary Fig. 1A): the spatial domain was circular with radius R = 1750 *μ*m and the temporal domain was exactly three days. Cells were seeded uniformly at the center of this domain along a radius *ρ*_0_ = 175 *μ*m at 36% confluence. To recapitulate the configuration of the MEMA profiling experiment, cells leaving the *ρ*_0_ domain can no longer adhere to the ECM and die. This was implemented by having the carrying capacity *K*(*x*) rapidly approach zero when *x* > *ρ*_0_. This setup can have energy attract cells to the periphery of a MEMA spot and beyond. We ran 161,000 simulations at variable energy consumption rates, chemotactic/haptotactic coefficients, energetic sensitivities, and diffusion rates of the growth-limiting resource (i.e. ECM-bound HGF; Table 1). 92% of these simulations completed successfully. For each simulation/ECM pair, we compared spatial distributions of in-silico and in-vitro confluence using the Wasserstein metric [23].

**Table 1.** Measured and inferred simulation parameters for one-population model (second column); and two-population model (third column). Parentheses indicate ranges with equally good fits. Corresponding non-dimensional parameters are shown in Supplementary Table 3. Net chemotaxis of cells is given by the interaction of the chemotactic coefficient (*χ*) and energy deficit inducing chemotaxis (*ξ_u_*), i.e. effective cell chemotaxis is approximately *χ*/*ξ_u_*. The larger *ξ_u_* the later the cells sense an energy deficit, i.e. the larger the energy gradient must be in order for them to accelerate their movement in response to it.

| Experimentally measured parameters | Symbol | Homogeneous | Heterogeneous | Unit |
| --- | --- | --- | --- | --- |
|  |  | cells on any ECM | cells |  |
| Maximal growth rate | $\lambda$ | 0.56 | 0.56 | day<sup>-1</sup> |
| Goer's seeding density | $u_0$ | 36.0 | 3.8 | % confluence |
| Grower's seeding density | $v_0$ | | 1.3 | % confluence |
| Simulation time | $t$ | 3 | 127 | days |
| Radius of domain | $\tilde{R}$ | 1750 | 1750 | $\mu\text{m}$ |
| Seeding radius | $\rho_0$ | 175 | 175 | $\mu\text{m}$ |
| Growth beyond $\rho_0$ | | no | yes | |

**Table 1.** Measured and inferred simulation parameters for one-population model (second column); and two-population model (third column).
| Inferred parameters | Symbol | Homogeneous | Heterogeneous | Unit |
| --- | --- | --- | --- | --- |
|  |  | cells on GAP43 | cells |  |
| Energy diffusion coefficient on ECM | $\Gamma_E$ | 9500 | 8982 | $\mu\text{m}^2 \text{ day}^{-1}$ |
| Energy consumption rate | $\delta$ | 57 | 23 | cells<sup>-1</sup> day<sup>-1</sup> |
| Energy needed for half-maximal growth | $\Phi_u$ | [0.01, 3] | 0.310 | |
| | $\Phi_v$ | | [0.05, 0.16] | |
| Chemotactic coefficient | $\chi$ | 5500 | 1919 | $\mu\text{m}^2 \text{ day}^{-1}$ |
| Energy deficit inducing chemotaxis | $\xi_u$ | [2, 300] | [1, 8] | |
| | $\xi_v$ | | 140 | |

The model was implemented in C++ (standard C++11). The armadillo package (ARMA version: 9.860.1) [24] was used for simulation of the PDEs. Simulations were run on a Intel Core i7 MacBook Pro, 2.6 GHz, 32 GB RAM. The source code is available at the github repository for the Integrated Mathematical Oncology department: GoOrGrow.

### Ploidy as biomarker of phenotypic divergence

We identified 44 breast cancer cell lines of known ploidy [25] and with available RNA-seq data in CCLE [26] and analyzed their drug sensitivity- and expression profiles as follows.

#### Drug sensitivity analysis

We used Growth rate inhibition (GR) metrics as proxies of differences in drug sensitivities between cell lines. Unlike traditional drug sensitivity metrics, like the *IC*_50_, GR curves account for unequal division rates, arising from biological variation or variable culture conditions – a major confounding factor of drug response [27]. Previously calculated GR curves and metrics were available for 41/44 breast cancer cell lines. A total of 46 drugs had been profiled on at least 80% of these cell lines and their *GR_AOC_* drug response metric [28] was downloaded from GRbrowser. For each drug we calculated the z-score of *GR_AOC_* across cell lines in order to compare drugs administered at different dose ranges. Of these 46 drugs, 39 could be broadly classified into two categories as either cytotoxic (25 drugs) or inhibitors of signaling pathways (14 drugs) (Supplementary Table 2). We then evaluated a cell line’s ploidy as a predictor of its *GR_AOC_* value using a linear regression model. Since molecular subtype of breast cancer cell lines is known to influence drug sensitivity we performed a multivariate analysis, including the molecular subtype as well as an interaction term between ploidy and drug category into the model.

#### RNA-Seq analysis

The molecular subtype classification of all cell lines was available from prior studies [29–32]. Of these 44 cases, four were suspension cell lines and excluded from further analysis. Of the remaining 40 cell lines, 20 originated from primary breast cancer tumors and were the focus of our analysis. Gene expression data was downloaded from CCLE. We used gene set variation analysis (GSVA) to model variation in pathway activity across cell lines [33]. Pathways for which less than ten gene members were expressed in a given cell lines were not quantified. The gene membership of 1,417 pathways was downloaded from the REACTOME database [34] (v63) for this purpose.

## Results

### High-ploidy breast cancer cell lines have increased metabolic activity and cell motility

To better understand the phenotypic profile of high-ploidy cells, we compared the ploidy of 41 breast cancer cell lines with their response to 46 drugs. As drug response metric, we used the integrated effect of the drug across a range of concentrations estimated from the ‘area over the curve’ (*GR_AOC_*) [27, 28]. We observed that cytotoxic drugs and drugs inhibiting signal transduction pathways were at opposite ends of the spectrum (Fig. 1A). Namely, ploidy was negatively correlated with the *GR_AOC_* for several cytotoxic drugs and positively correlated with the *GR_AOC_* of various mTOR inhibitors, suggesting high ploidy breast cancer cell lines tend to be resistant to DNA damaging agents, while sensitive to drugs targeting nutrient sensing and motility.

**Figure 1.**
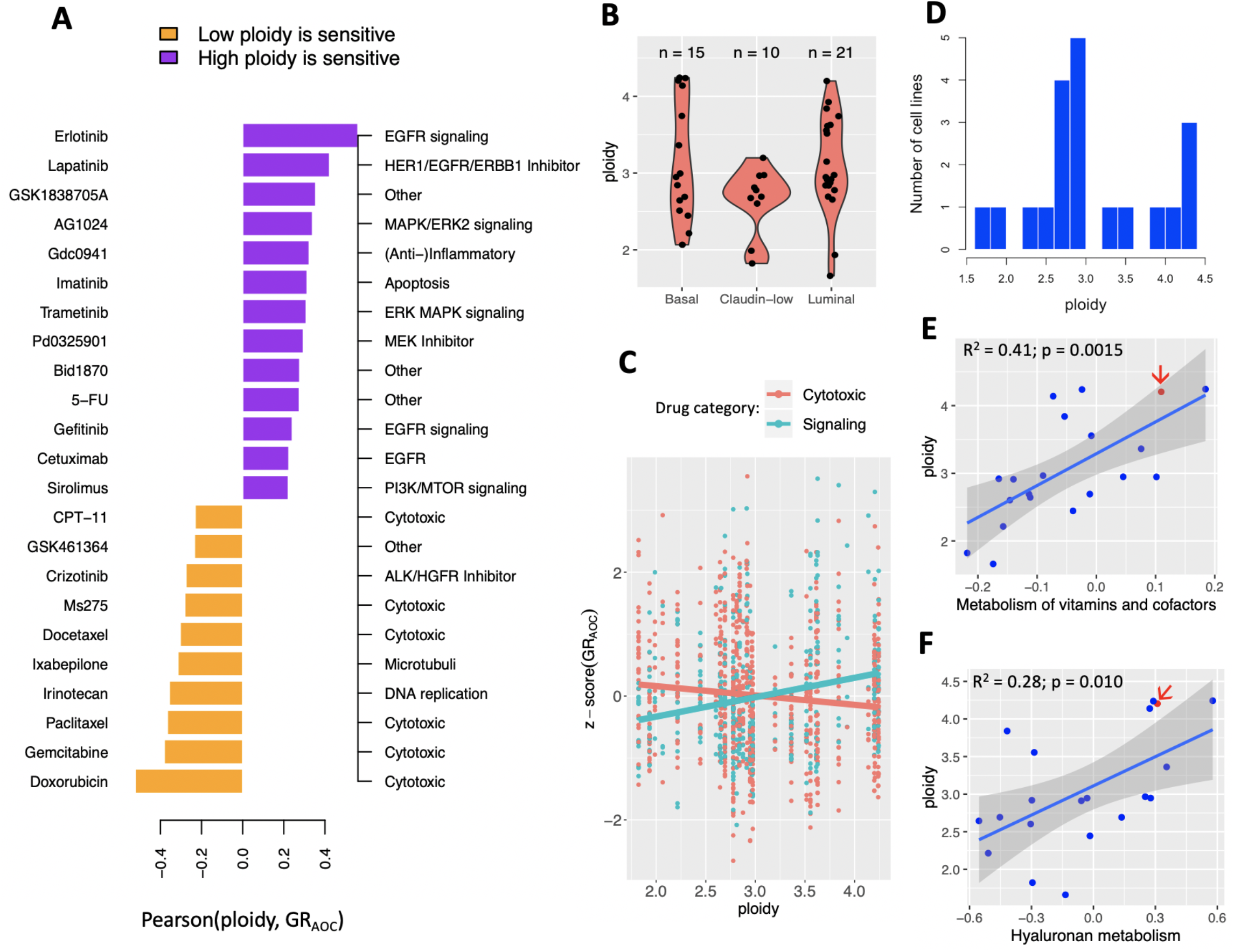
Ploidy, pathway activity and drug sensitivity across breast cancer cell lines from CCLE. (A) High-ploidy breast cancer cell lines are resistant to cytotoxic drugs, but tend to be more sensitive to inhibitors of mTOR, EGFR and MAPK signaling pathways. Hereby ploidy is defined as the number of chromosomes in the cell line’s consensus karyotype, weighted by chromosome size. Only drugs with a Pearson correlation coefficient at or above 0.2 are shown here. (B) Distribution of ploidy within and across three molecular breast cancer subtypes. (C) Regression coefficient of ploidy as predictor of *GR_AOC_* has opposite signs depending on drug category across all subtypes. (D) Distribution of ploidy across 20 primary, adherent breast cancer cell lines from CCLE. (E-F) Ploidy is correlated with the activity of pathways involved in metabolism of vitamins and cofactors (E) and Hyaluronan metabolism (F). One cell line with available MEMA profiling data – HCC1954 – is highlighted (red arrow).

We built a multivariate regression model of drug sensitivities to test the hypothesis that the relationship between ploidy and *GR_AOC_* was different for cytotoxic drugs than for inhibitors of cell signaling pathways. Molecular subtype alone (Fig. 1B), could explain 0.4% of the variability in *GR_AOC_* z-scores across cell lines (adjusted R-square = 0.0044; p = 0.026). Including ploidy into the model did not improve its predictive accuracy (adjusted R-square = 0.0037; p = 0.058). However, an interaction term between ploidy and drug category (cytotoxic: 27 drugs vs. signaling: 16 drugs) increased accuracy to explain 2.6% of variability in drug sensitivity across cell lines (adjusted R-square = 0.026; p < 1e-5; Fig. 1C). The same improvement from an interaction term between ploidy and drug category was observed in an independent dataset of maximal inhibitory concentration (*IC*_50_) values of 34 cytotoxic drugs and 51 signaling inhibitors obtained from the GDSC (Genomics of Drug Sensitivity in Cancer) database [35] (Supplementary Fig. 3).

We then focused on a subset of aforementioned 41 cell lines, namely those that had been established from primary breast cancer tumors as adherent cells (20 cell lines; Fig. 1D) and we quantified their pathway activity (see Methods). A total of 27 pathways were correlated to ploidy at a significant p-value (| Pearson r | ≥ 0.44; p ≤ 0.05; Supplementary Table 1). The strongest correlations were observed for metabolic pathways such as hyaluronan metabolism and metabolism of vitamins (Fig. 1E-F). Hyaluronic acid is a main component of the ECM and its synthesis has been shown to associate with cell migration [36, 37].

These results support a model that connects high ploidy with both, the chemotactic ability and metabolic energy deficit of a cell.

### Infiltration of homogeneous populations

Model design was guided by the goal to describe growth dynamics along two axes: from random to directed cell motility and from homogeneous to heterogeneous cell compositions (Methods equations (2a) - (2c)). While the last section will step into the second axis, the following two subsections distinguish scenarios along the first axis: (i) “homogeneous nutrient environments” are environments in which random cell motility dominates throughout population growth; (ii) “heterogeneous nutrient environments” imply the formation of gradients which cause cells to move in a directed fashion, as is the case during cellular growth on an ECM.

#### Homogeneous nutrient environments

When the diffusion of the nutrient occurs on a much faster time scale than the actions taken by cells, we can assume that at the time scale of cells, the nutrient is essentially uniform in space. This simplification allows us to neglect chemotactic/haptotactic motion and consider only random cell motility as the driving force that spreads the cell density throughout the dish.

We obtain analytical estimates for the degree of infiltration in a homogeneous environment that lays the groundwork for new predictions. To arrive at (3a)-(3b), we employed the approximations that diffusion of energy molecules (e.g. glucose) is fast relative to cell movement and that the cells’ movement through the dish can be approximated by a traveling wave [38]. These assumptions were verified by comparing the front estimates with results from the full numerical model (equations (2a) - (2c); Fig. 2A,B).

**Figure 2.**
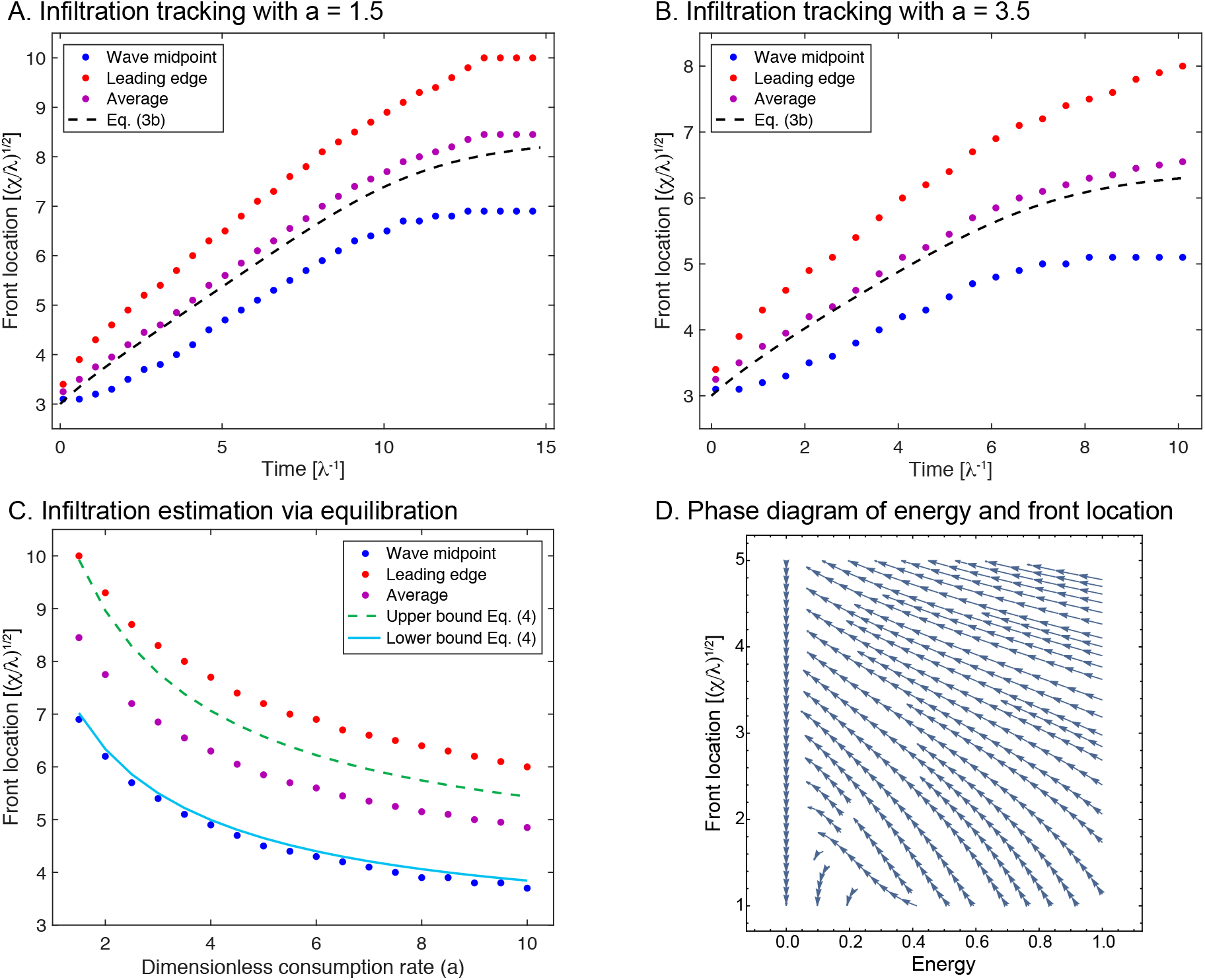
Comparing analytical approximations of the degree of infiltration with those obtained from simulations. (A-B) Traveling-wave solutions at energy consumption rates *a* = 1.5 (A) and *a* = 3.5 (B). (C) Upper and lower boundaries of traveling wave solutions estimated from equilibration are shown as a function of consumption rate. Approximation is found by lower and upper bound Λ = 0, 1 from equation (4). (D) Phase diagram of energy consumption and front location using the derived coupled system (3a)-(3b). (A-D) All approximations and simulations assume energy is uniformly distributed at all times, i.e. chemotaxis does not take place. Parameter values for initial seeding radius (*ρ*_0_), dish radius (*R*), and sensitivity to low energy (*ϕ*) are set to 3, 10 and 0.05 respectively. Red = leading edge of wave (estimated by finding value of cell concentration closest to 0.01); blue = mid point of wave (estimated by finding value of cell concentration closest to 0.5); purple = average of red and blue; black lines are approximations based on analytical solutions. Time is in units of the maximal growth rate of the given cell line. Front location is given in units of the characteristic lengthy 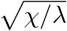.

An interesting alternative to tracking the wave over time is to assume it travels as a wave, but only record the density after the system has reached uniformity. If the death rate is much smaller than the time needed for the cells to spread uniformly, we can bound the degree of infiltration that occurred from only knowing the uniform density of cells (eq. (4); Fig. 2C).

These analytical solutions point to scaling relationships for the speed of the moving front. For highly efficient energy-using cell lines (*ϕ* ≪ 1) (Fig. 2D), the front will evolve at a speed nearly independent of energy. In contrast, for large *ϕ* ≫ 1, the speed of the front falls off as 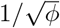. These predictions of the behavior of infiltration on parameters can be investigated experimentally.

#### Heterogeneous nutrient environments

Assumptions made in the prior section apply to standard cell cultures of adhesive cells in a typical cell culture dish, where energetic resources diffuse so fast that gradients do not form. These assumptions break down during cellular growth on an ECM. Binding to the ECM can cause soluble factors (like HGF) to act and signal as solid-phase ligands [39, 40]. Proteolytic degradation of these ECMs then creates haptotactic gradients. Fig. 1 includes HCC1954 – a near-tetraploid breast cancer cell line whose growth on various ECMs has been measured via MEMA profiling. We analyzed HCC1954 and looked to determine if our mathematical model can explain its spatial growth patterns. MEMA profiling resulted in considerable variability of growth patterns across different ECMs (Fig. 3A,B).

**Figure 3.**
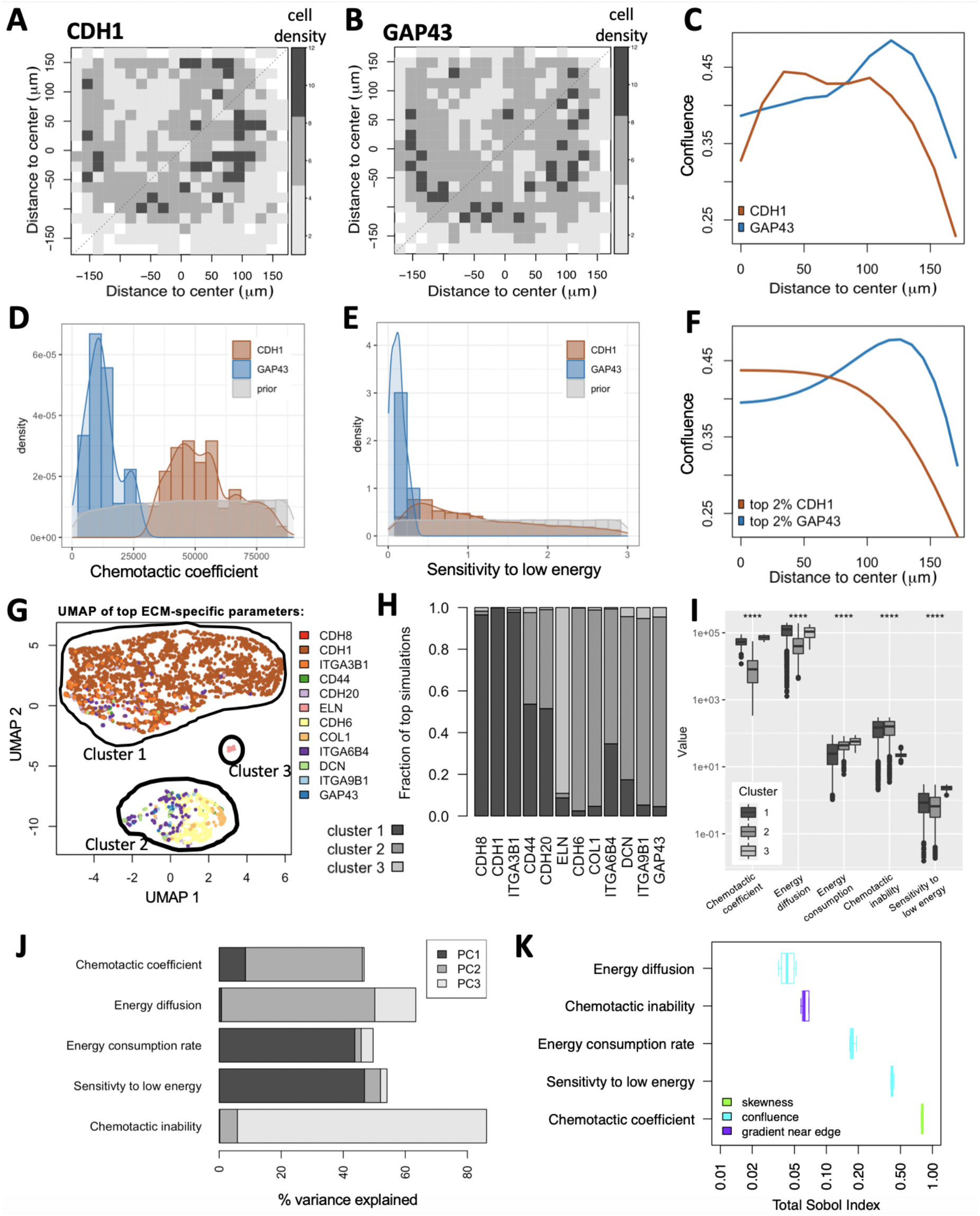
Model calibration using MEMA profiling of HCC1954 cells. (**A-C**) **Experimentally measured data**. Variability in cell growth patterns across ECMs is demonstrated via two example ECMs: CDH1 (A) and GAP43 (B). The average local cell densities are displayed for both (color legend). (C) Projecting their 2D spatial distributions onto 1D reveals enrichment of cells at the edge of the ECM spot for GAP43, but not for CDH1. (**D-F**) **Simulated data**. Comparing prior-distribution of chemotactic coefficients (D) and sensitivity to low energy (E) to ECM-specific posterior distributions reveals clear differences between CDH1 and GAP43 for both parameters. (F) Maximum-likelihood parameter choices for CDH1 and GAP43 result in distinct spatial distributions between the two ECMs, each of which resemble the measured distributions (C). (**G-I**) **ECM specific model parameters**. Simulations were compared to each measured ECM-specific growth pattern and ranked by their maximum similarity. (G-H) The five model parameters from the top 2.3% simulations were projected onto UMAP space, revealing three clusters. Color-coding simulations by the ECM responsible for their presence in the top simulations suggests enrichment of most ECMs to only a single cluster (G). (H) This was confirmed when comparing cluster membership across the 12 represented ECMs. (I) All five model parameters (x-axis) show significant differences between the three clusters (****: p <= 0.0001). (**J-K**) **Parameter sensitivity analysis**. (J) The % variance explained per parameter shows significant contribution of all five model parameters to at least one of the first three principal components (PCs). (K) The Sobol index (x-axis) tells us which parameter best explains which aspect of the cells’ spatial distribution (color-code): its skewness, confluence or gradient near the edge.

We projected 2D spatial distributions measured on the MEMA array onto one dimension (Fig. 3C), rendering them comparable to those obtained from our simulations (Fig. 3D-E). For each simulation/ECM pair, we calculated the distance between in-silico and in-vitro spatial cell distributions using the Wasserstein metric (Fig. 3C,F) and ranked simulations by their minimum distance across ECMs. The top 2.3% simulation parameters were then stratified by the ECM whose spatial pattern they best resemble and compared to uniform prior parameter distributions (Fig. 3D,E). 75% of the ECMs accounted for only 1.55% of the top simulations. The tendency of cells on these ECMs to grow in a toroidal shape was strong, suggesting it may be the consequence of non-uniform printing of ECMs onto the array. We concluded that our model cannot explain the growth patterns on those ECMs well and focused our attention on the remaining 12 ECMs, represented by 3,429 simulations. We refer to the parameters of these simulations as inferred parameter space. Principal component analysis (PCA), Uniform Manifold Approximation and Projection (UMAP) [41] and density clustering [42] of the inferred parameter space revealed three clusters (Fig. 3G), with different ECMs segregating mainly into different clusters (Fig. 3H).

The two largest clusters differed mostly in their chemotactic/haptotactic- and energy diffusion coefficients; while the small cluster stood out by a high sensitivity to low energy and fast chemotactic/haptotactic response (Fig. 3I). Overall, all five model parameters showed significant differences between the three clusters, suggesting they all contribute to distinction between ECM growth patterns (Fig. 3I). This was further affirmed when looking at the % variance explained per principal component per parameter (Fig. 3J). To formalize parameter sensitivity analysis independent of ECMs, we also calculated the Sobol index [43] of each parameter. The Sobol Index quantifies how much of the variability in spatial cell concentration is explained by each parameter, while accounting for all its interaction effects. Each parameter contributed to significant variance (Sobol Index > 0.02; [43]) in at least one of three spatial statistics (Fig. 3K): skewness, confluence and gradient near the edge of the ECM.

We observed substantial differences in chemotactic/haptotactic coefficients and energy consumption rates between ECMs (Supplementary Fig. 4). To query the biological significance of this variability we quantified the expression of the 12 ECMs in the HCC1954 cell line (Methods). Two of the five inferred ECM-specific model parameters were correlated with RNA-seq derived expression of the corresponding ECM: energy consumption rate (Pearson r = −0.657, p = 0.028) and sensitivity to low energy (Pearson r = 0.562, p = 0.071) – though latter fell short above significance (Supplementary Fig. 5).

In summary, the posterior distributions of model parameters represent a substantial departure from the uniform priors and could explain a significant proportion of growth conditions on the HGF-exposed MEMA array. This approach identified regions of interest in the parameter search space, allowing us to focus further simulations on biologically relevant chemotactic/haptotactic coefficients and energy diffusion rates.

### Infiltration of heterogeneous, chemotactic populations

Growth of cells in a given ECM environment was measured across 13-30 replicates on the MEMA platform. While our model – when calibrated to the corresponding ECM environment – could explain the observed growth pattern in the majority of these replicates, a substantial fraction could not be explained by fixed choices of sensitivity to low energy and directed cell motility (Supplementary Fig. 4). One possibility that may explain this is that HCC1954 is a heterogeneous cell line, with clones of variable phenotypes co-evolving. Representation of these clones among the 31 cells that were on an average sampled for each replicate may vary (Supplementary Fig. 2). This hypothesis is supported by a bimodal distribution of DNA content observed among replicating HCC1954 cells on individual ECM spots (Fig. 4A,B). If the HCC1954 population was homogeneous, we would expect a unimodal distribution of DAPI intensity among S-phase cells of this cell line. The observation of a bimodal distribution among S-phase cells suggest that HCC1954 is likely a polyploid cell line, i.e. clones of variable ploidies co-exist in this cell line.

**Figure 4.**
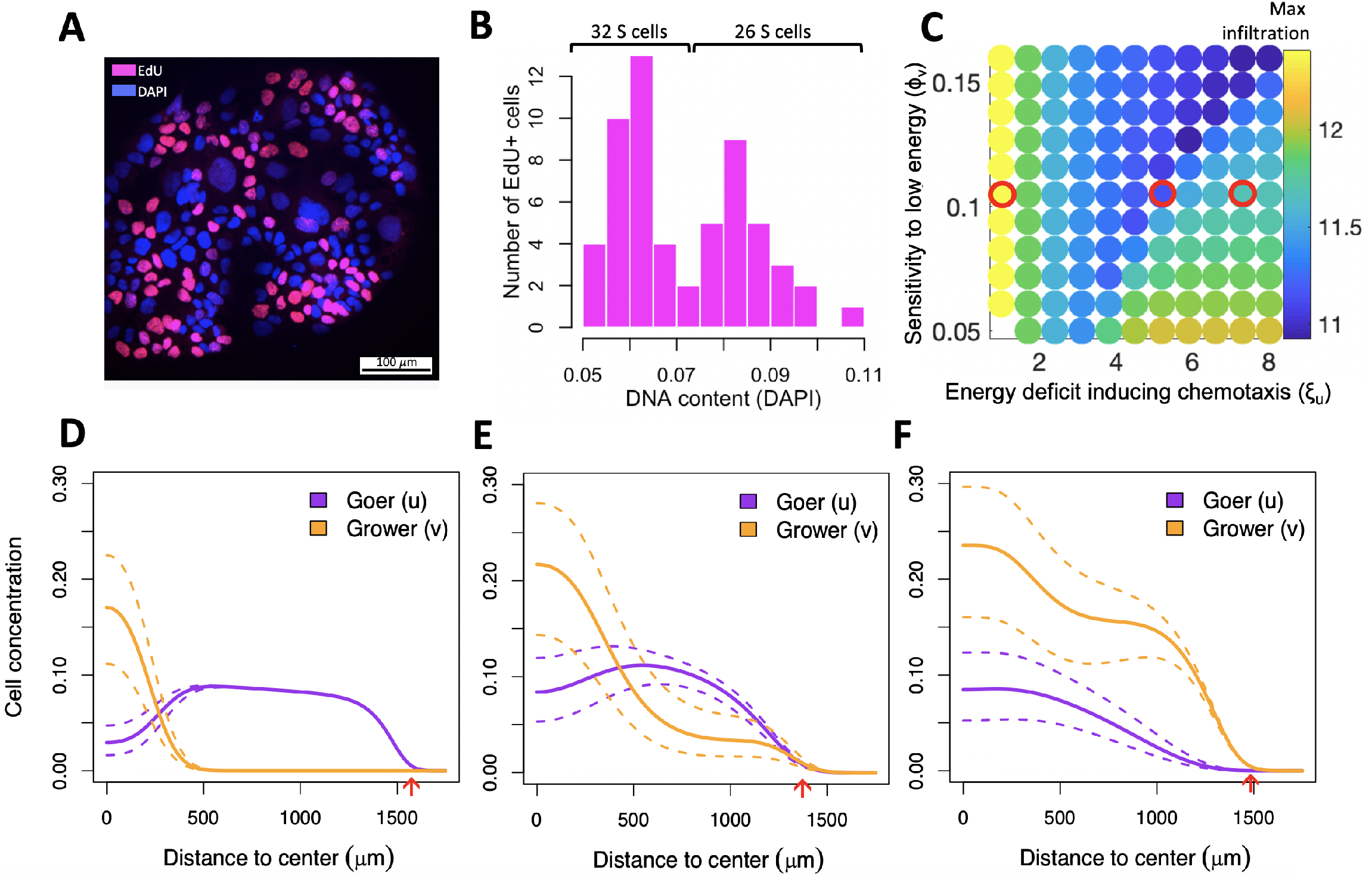
Internal competition of co-existing subpopulations for same space slows down invasion of the metapopulation. (A) DNA content and cell cycle state of 162 cells growing on HGF-exposed ICAM1. (B) DAPI intensity of 58 replicating (EdU+) cells shows abimodal distribution, indicating the presence of two subpopulations – a low-ploidy population (grower) comprising 55% cells and a high-ploidy population (goer) comprising 45% cells. (C) Arms race between the grower’s energetic sensitivity (y-axis) and the goer’s chemotactic ability (x-axis) reduces infiltration distance (color bar). Red circles outline parameter combinations of interest explored in (D-F). (D-F) Spatial distribution of goer and grower for parameter values outlined in (C). Dotted lines outline extreme trajectories of expected cell concentrations due to incertitude in initial goer/grower proportions, as estimated from the silhouette coefficient of cells in panel B (see also Supplementary Fig. 6B). (D) High chemotactic motility will cause the goer to leave the center of the dish too soon, leaving room for the grower to expand there. (E) With an intermediate motility the goer succeeds maintaining high representation both at the center and edge of the dish. (F) Low motility will prevent the goer from gaining a sufficient spatial lead from the grower while energy is still abundant, and it will lose dominance at the edge of the dish once energy becomes sparse. Red arrows indicate maximum infiltration distance achieved by either of the two populations.

To better understand the growth dynamics in a polyploid population, we used the two-subpopulation version of our model, whereby variable chemotactic abilities and energetic sensitivities of goer- and grower subpopulations compete with one another (equations (2a) - (2c)). We used fixed values for energy diffusion- and consumption rates as informed by model calibration (Fig. 3B) and varied sensitivity to low energy and chemotactic ability of both goer and grower, subject to equations (2a) - (2c) (Table 1). We initially used the same spatial and temporal domains as during model calibration, but concluded that the implied duration of the experiment (3 days) was too short for dynamics between the two populations to manifest. Each MEMA spot has a low capacity, whereby confluence is reached at no more than a few hundreds of cells. Such a small number of cells will not exhibit wave-like behavior and therefore will not suffice for spatial structure to emerge. We therefor extended temporal and spatial domains of our simulations, seeding cells at a lower confluence and letting them grow onto the entire energy domain until they consume all available energy (average of 127 days; Table 1).

We observed a non-monotonic relation between the goer’s chemotactic ability and the speed with which the metapopulation invades the dish, with intermediate values being the least beneficial to its growth and spread (Fig. 4C). Temporal analysis of the simulations (Supplementary Data 1-3), revealed that if the goer’s chemotactic motility is too high, it will leave the center of the dish too soon, leaving room for the grower to expand locally (Fig. 4D). By contrast, if the goer’s motility is too low, it will miss the time-window of opportunity to ensure its dominance further away from the center of the dish while energy is still abundant. As a consequence, it will be outgrown by the grower at the edge of the dish once energy becomes sparse (Fig. 4E). Only when the goer has an intermediate motility, does the grower persistently coexist with it, both at the center and edge of the dish (Fig. 4F).

## Discussion

Models of infiltration are typically formulated under two critical assumptions. First, that energy production and consumption are non-uniform, leading to the formation of an energy gradient [44–46]; or second, that energy consumption is very slow compared to production, leading to an essentially infinite energetic resource [47]. Here we formulate a generalized model of infiltration when energy is finite and investigate its behavior along a spectrum of scenarios, from permanent energy uniformity to scenarios where this uniformity is gradually lost. The model derivation does not assume a particular dimension (e.g. 2D *in vitro* experiments vs. *in vivo* or 3D spheroid experiments). Many parameters that were valid in 2D will also extend naturally to 3D. For example, we would not expect a difference in consumption rates or half-maximal growth rates of the cells. However, energy diffusion (Γ_E_) or random cell motion (Γ) will be higher due to the increased degree of freedom [48]. When energy is uniformly distributed at all times and the time scale for cell death is substantially longer than that of cell motility and birth, our results suggests that the degree of infiltration can be approximated using the cells’ density at equilibration of movement and growth (Fig. 2C).

With an energy gradient that becomes steeper over time, our analytical approximations no longer hold, as directed cell movement becomes non-negligible. For this scenario, we leveraged MEMA profiling to inform regions of interest in the parameter search space. These regions of interest are relevant for cellular growth on a variety of HGF exposed ECM proteins. We observe correlations between inferred model parameters and RNA-seq derived signatures, even though the latter were not used during parameter inference. A potential explanation for the negative correlation between ECM-specific energy consumption and expression is that our model does not account for the possibility that cells can replace the ECM they degrade. The slower the rate of this replacement is, the higher the consumption rate appears to be. On the other hand, the more dependent cells are on a ECM for growth, the faster they must replace it, potentially explaining a positive correlation between ECM-specific expression and sensitivity to low energy.

When calibrating our model to a given ECM environment, growth patterns of a substantial fraction of replicates of that ECM could not be explained by fixed choices of sensitivity to low energy (Supplementary Fig. 4). A potential explanation for this are variable cell compositions across experimental replicates. An alternative explanation is that this variability stems from artifacts that arise during non-uniform printing of ECMs onto the array–the so called ring effect. However, a bimodal distribution was also observed in the DNA content of replicating cells, which is not affected by potential printing artifacts. The second peak of this bimodal distribution was wider, consistent with the fact that high-ploidy cells with more DNA need longer to replicate.

The cell line HCC1954 is described as a hyper-tetraploid cell line with an average DNA-content of 4.2 [25]. However, this average value may be misleading, as suggested by stark variability in nuclei sizes (Fig. 4A). Despite a wealth of genomic information generated for this cell line [25], to the best of our knowledge no prior reports indicate whether or not the cell line is polyploid. We and others have found that high ploidy is an aneuploidy-tolerating state that accompanies intra-tumor heterogeneity in vivo and in *vitro* [5, 49, 50]. Our results suggest that HCC1954 is likely polyploid.

One event that could have led to this polyploid state is WGD. In contrast to cell lines, WGD events in primary tumors are mostly clonal, not subclonal [5, 6, 10] – clones carrying a doubled genome often sweep over the population, such that by the time the tumor is detected, the diploid ancestor no longer exists. A related scenario are advanced, therapy-exposed tumors shown to revert back to genomic stability, potentially bringing a WGD population back to a genomic state that more closely resembles its diploid ancestral state [51]. The model presented here can investigate how dynamics between the two subpopulations unfold in both of these scenarios – early, shortly after the WGD or late, after therapy exposure. This would characterize what circumstances prevent the WGD carrying clone from becoming dominant or from retaining its dominance and could help explain WGD incidence in primary and recurrent tumors.

If spatial and temporal domains were to be extended beyond the configuration of MEMA spots, our simulations predict that spatial segregation of two co-existing subpopulations according to their ploidy is a likely scenario and depends on the energy consumption rate. Our model can easily be extended to more than two subpopulations, for example to include a subpopulation of normal cells. For each additional cell type, a new compartment can be added to the model, with growth- and motion-related parameters that are specific to the corresponding cell type. However, the model currently does not incorporate mutations, i.e. the process of generating new clonal lines. A next step will be to extend our model to include mutation events, specifically chromosome mis-segregations that contribute extensively to diversify ploidy of a population [52, 53]. The additional DNA content of high-ploidy cells, though energetically costly, brings a masking effect against the deleterious consequences of chromosome losses [10]. This duality may explain the higher sensitivity to glycolysis inhibitors of high-ploidy cells and their lower sensitivity to cytotoxic drugs previously reported in Glioblastoma [54].

In line with prior reports we find that increased resistance of breast cancer cell lines to cytotoxic drugs is associated with high ploidy. In contrast, high ploidy breast cancer cell lines were sensitive to inhibitors of signal transduction pathways, including EGFR and especially MTOR signalling. A commonality among those pathways is their contribution to a cell’s chemotactic response [55–57], suggesting opportunities to tune chemotaxis. MTOR inhibitors (mTOR-I), such as Rapamycin, significantly decrease migration of breast cancer cells in a dose-dependent manner [58, 59]. Rapamycin inhibits cell motility by a mechanism similar to that by which it inhibits cell proliferation [60], suggesting that the mTOR pathway lies at the intersection of a cell’s decision between proliferation and migration. If high ploidy is indeed a characteristic specific to goer-like cells, then mTOR-Is are likely affecting this cell type (Fig. 1A,E,F), and could be used to inhibit its chemotactic response, thereby moving the population up the *x*-axis of Fig. 4C. Delaying chemotactic response of highly chemotactic cells could slow down invasion by maximizing competition within a polyploid population. If on the other hand chemotactic response of high ploidy cells is already at an intermediate level, our simulation suggest that further reduction may accelerate invasion of low ploidy cells. For such scenarios therapeutic strategies that include an mTOR-I may not be successful. Experiments will be needed to verify these in-silico results in-vitro. Knowing how co-existing clones with differential drug sensitivities segregate spatially can offer opportunities to administer these drug combinations more effectively.

## Supporting information

Simulated Data Fig 4D

Simulated Data Fig 4F

Simulated Data Fig 4E

## Acknowledgements

We thank the researchers in the Integrated Mathematical Oncology department at Moffitt Cancer Center for fruitful discussions and early feedback on parts of the manuscript. The content is solely the responsibility of the authors and does not necessarily represent the official views of the National Institutes of Health or the H. Lee Moffitt Cancer Center and Research Institute.

