## Supplementary methods, figures and tables for "Invasion of homogeneous and polyploid populations in nutrient-limiting environments"

**Gregory J. Kimmel[1], Mark Dane[2], Laura Heiser[2,1], Philipp M. Altrock[1] and Noemi Andor[1*]**
*[1]Department of Integrated Mathematical Oncology, Moffitt Cancer Center, Tampa, FL, USA [2]Department of Biomedical Engineering, OHSU Center for Spatial Systems Biomedicine, Knight Cancer Institute, Oregon Health & Sciences University, Portland, OR, USA*

#### Financial support

This work was supported by the National Cancer Institute R00CA215256 awarded to NA. PMA acknowledges support through the National Cancer Institute, part of the National Institutes of Health (NIH), under grant number P30-CA076292. LH acknowledges support through NIH research grants 1U54CA209988, U54-HG008100, Jayne Koskinas Ted Giovanis Foundation for Health and Policy, and Breast Cancer Research Foundation.

#### Conflict of Interest

The authors declare no potential conflicts of interest.

#### Keywords

Evolutionary tradeoffs, Aneuploidy, Polyploidy, Breast cancer, Mathematical Modeling, Whole genome doubling, Partial differential equations, Traveling waves

[*]Noemi Andor; Moffitt Cancer Center, SRB 23224 B2 3011 Holly Drive, 33612 Tampa;

### Supplementary Information

#### Model derivation

We suppose that the energy flux is driven entirely via diffusion with coefficient $\Gamma_E$ dependent on the type of media. The energy is consumed in proportion to the amount of cells present. For cell motility, we assume it is driven both by chemokinesis and chemotaxis. This leads to our general coupled system

$$\frac{\partial \tilde{E}}{\partial t} = \Gamma_E \nabla^2 \tilde{E} - \delta \tilde{E}(U+V), \tag{S1a}$$

$$\frac{\partial U}{\partial t} = \nabla \cdot \left( \Gamma \nabla U - \chi U \frac{\nabla \tilde{E}}{\Xi_U + \tilde{E}} \right) + \lambda \frac{\tilde{E}}{\Phi_U + \tilde{E}} U \left(1 - \frac{U+V}{K}\right), \tag{S1b}$$

$$\frac{\partial V}{\partial t} = \nabla \cdot \left( \Gamma \nabla V - \chi V \frac{\nabla \tilde{E}}{\Xi_V + \tilde{E}} \right) + \lambda \frac{\tilde{E}}{\Phi_V + \tilde{E}} V \left(1 - \frac{U+V}{K}\right). \tag{S1c}$$

The system (S1) is defined on a dish of radius $R$ subject to no-flux boundary conditions. We assume the cells $U, V$ are initially concentrated at the center with radius $\rho_0 \leq R$ and initial concentrations $U_0, V_0$. Further, we assume that the energy density is uniformly distributed on the plate with initial value $E_0$. All parameters except otherwise stated are independent of any of the state variables $(\tilde{E}, U, V)$. The energy is consumed at rate $\delta$. Both cells can divide at the maximal rate $\lambda$, but are restricted by the energy density $\tilde{E}$. The cells can grow to a local maximal density given by $K$. This parameter is often cell line-dependent and is related to contact inhibition and cells being able to grow on top of each other.

Before proceeding, we rescale our variables $\tilde{E}, U, V$. Define $E = \tilde{E}/E_0, u = U/K, v = V/K$. In other words, we are looking at the change in energy relative to the initial amount and normalizing the cells by their local carrying capacity density. This leads to a natural redefinition of the energy sensitivity parameters $\xi_i = \Xi_i/E_0$ and $\phi_i = \Phi_i/E_0$.

Next, we rescale time in terms of the maximal growth rate $\tau = \lambda t$ and introduce a characteristic length by $L = \sqrt{\chi/\lambda}$. With this rescaling, we have our main (non-dimensional) equations

$$\frac{\partial E}{\partial \tau} = \gamma_E \nabla^2 E - aE(u+v), \tag{S2a}$$

$$\frac{\partial u}{\partial \tau} = \nabla \cdot \left( b\nabla u - u \frac{\nabla E}{\xi_U + E} \right) + \frac{E}{\phi_U + E} u(1-u-v), \tag{S2b}$$

$$\frac{\partial v}{\partial \tau} = \nabla \cdot \left( b\nabla v - v \frac{\nabla E}{\xi_V + E} \right) + \frac{E}{\phi_V + E} v(1-u-v), \tag{S2c}$$

where we have defined $\gamma_E = \Gamma_E/\chi,\ a = \delta K/\lambda,\ b = \Gamma/\chi$.

We made a few choices that warrant further explanation. First, we were mainly interested in the degree of infiltration that the heterogeneous population would make given a finite source of energy. If the source was infinite, the growth rate term would never decay, and the cells would eventually make their way across the entire dish (both in simulations and eventually in experiments through random motion, if permitted). This energy source term is also nontrivial. For example, we can use a periodic energy source term where $q = q_0 \sin(2\pi t/T)$, or we can use a constant source term $q = q_0$. Ultimately, any source term which does not decay to zero at long times will eventually have the cells occupying the entire dish. Since we are mostly interested in how far they can move given a finite amount of energy, we avoided adding this additional term at this time.

Second, the inclusion of a local carrying capacity is to prohibit overgrowth of too many cells on top of each other. Different cell lines are expected to have different $K$'s and cells that don't have this restriction can be modeled by taking $K \to \infty$. This also avoids potential overflow, when the half-maximal energy parameters $\phi \ll 1$. Ultimately, energy-dependent growth in a finite resource environment is a natural mechanism that stops growth, but a local carrying capacity is also a natural parameter as we don't expect unbounded stacking of cells.

Third, additional compartments that model the tumor microenvironment (e.g. normal cells, matrix tissue) were purposely omitted. The model has the ability to incorporate this through increasing complexity in a modeling hierarchy by making the parameters related to growth and motion dependent on these additional compartments. For immobile structures such as matrix tissue, we could generalize parameters to be spatially-dependent. For additional cell types (e.g. normal cells), we can add compartments to the model with growth- and motion-related parameters that are specific to the corresponding cell type. In this regard, our model is flexible and amendable to further additions.

#### Infiltration

An important question in cancer dynamics is infiltration into and through tissue. The desire for cells to move is inherently tied to the availability of nutrients and space. To this end we define $\Psi(\tau) := [\rho(\tau) - \rho_0]/\rho_0$ where $\rho(0) = \rho_0$, the initial radius of cell seeding density. $\Psi$ can be thought of as a non-dimensional measure of infiltration attained after time $\tau$. This dimensionless measure has the added benefit of being scale-independent. An inherent difficulty with random cell motility and calculating infiltration is that the system always reaches the boundary in finite time. Instead we will define the maximum degree of infiltration to be given by the time needed for the total energy to be below a threshold $\varepsilon \ll 1$. For simulations, we took the 1-norm $\|E\|_1 < \varepsilon = 1e-4$.

In general, the maximum degree of infiltration is difficult to predict analytically, so we will only consider the single population case when obtaining our analytical estimates. We will also make use of the simplification that most energy-type molecules (e.g. glucose) have a diffusion coefficient that is very large, relative to cell movement. This allows us to write the reduced model:

$$\frac{dE}{d\tau} = -a \left(\frac{\rho}{R}\right)^2 E(u+v), \tag{S3a}$$

$$\frac{\partial u}{\partial \tau} = b\nabla^2 u + \frac{E}{\phi_u + E} u(1-u), \tag{S3b}$$

The additional term $(\rho/R)^2$ arises due to the fact that energy is now assumed to be homogeneous in space. Hence, (S2a) can be integrated over space, but we must account for the fact that $E$ is uniform and $u$ is non-zero only if $r \leq \rho(\tau)$. Performing this integration leads to the modification in (S3a).

**No chemotaxis infiltration estimate**

Consider (S3a) - (S3b), we write the radially symmetric Laplacian explicitly and obtain

$$\frac{dE}{d\tau} = -a\left(\frac{\rho}{R}\right)^2 Eu, \tag{S4}$$

$$\frac{\partial u}{\partial \tau} = b\left(\frac{\partial^2 u}{\partial r^2} + \frac{n-1}{r}\frac{\partial u}{\partial r}\right) + \frac{E}{\phi + E}u(1-u), \tag{S5}$$

where $n$ is the dimension (typically $n = 2, 3$, for circular or spherical growth, respectively). We now seek traveling wave-like solutions, where we assume that the wave speed is a slow function of $r$ and $\tau$. The solution obtained will verify that these assumptions are valid for our system. Our ansatz takes the form $u(r, \tau) = U(r - \eta\tau) = U(z)$. Note that in spatial equilibrium, $u = 1$ is stable and $u = 0$ is an unstable steady state. If the unstable state is what governs the wave speed, then the wave is said to be "pulled", otherwise it is "pushed". Based on numerical simulations, we assume that the state $u = 1$ is traveling across the domain and therefore set $u = 1$ in (S4):

$$\frac{dE}{d\tau} = -a\left(\frac{\rho}{R}\right)^2 E, \tag{S6}$$

$$0 = b\frac{\partial^2 U}{\partial z^2} + \left(\eta + \frac{b(n-1)}{r}\right)\frac{\partial U}{\partial z} + \frac{E}{\phi + E}U(1-U). \tag{S7}$$

Following a similar analysis conducted by Kaper et. al. [59], we define the new spatial variable $r = z + \eta t = \rho + \tilde{z}$. Treating $\rho$ and $\eta$ as slowly varying functions we can replace the above equation in terms of $\tilde{z}$ ,

$$\frac{dE}{d\tau} = -a\left(\frac{\rho}{R}\right)^2 E, \tag{S8}$$

$$0 = b\frac{\partial^2 U}{\partial \tilde{z}^2} + \left(\eta + \frac{b(n-1)}{\rho + \tilde{z}}\right)\frac{\partial U}{\partial \tilde{z}} + \frac{E}{\phi + E}U(1-U). \tag{S9}$$

Treating $\rho$ as a large constant gives the effective speed:

$$\tilde{\eta} = \eta + \frac{b(n-1)}{\rho}. \tag{S10}$$

We identify $\eta = d\rho/d\tau$ (the speed of the moving front) and so we have the evolution of the front given by the following coupled system:

$$\frac{dE}{d\tau} = -a\left(\frac{\rho}{R}\right)^2 E, \tag{S11}$$

$$\frac{d\rho}{d\tau} = 2\sqrt{\frac{bE}{\phi + E}} - \frac{b(n-1)}{\rho}. \tag{S12}$$

We see that our assumptions on the behavior of the traveling wave are verified. Since $\rho$ is assumed much larger than 0 and $\rho/R \ll 1$, $E$ is a slowly varying function of time and $\rho$ is a slowly varying function of time and its size.

**Estimating the degree of infiltration from equilibration**

An interesting alternative to tracking the wave over time is to assume it travels as a wave, but only record the density after the system has reached uniformity. This is possible if the death rate (which has been neglected) is much smaller than the time it would take the cells to spread uniformly. If this is the case, we can bound the degree of infiltration from only knowing the uniform value at the end of the experiment. To see this, note that at the end of the wave, with energy now 0, the total number of cells $u_T$ should remain fixed and so:

$$u_T = 2\pi \int_0^{\rho(T)} ur\, dr = 2\pi\rho(T)^2 \int_0^1 u(\rho s)s\, ds. \tag{S13}$$

The uniform value $\bar{u}$ at confluence leads to:

$$u_T = 2\pi\bar{u}\int_0^R r\, dr = \pi\bar{u}R^2. \tag{S14}$$

Equating these gives an equation for the unknown $\rho(T)$:

$$[\rho(T)]^2 = \frac{\bar{u}R^2}{2\int_0^1 u(\rho s)s\, ds}. \tag{S15}$$

Note that $s \in [0, 1]$ is a measure of distance in terms of the location of the front. Let $\Lambda$ be the width of the transition zone from $u = 1$ to $u = 0$, then we may partition the integral noting that $s < \Lambda$ implies that $u \approx 1$

$$\int_0^1 us\, ds = \int_0^\Lambda us\, ds + \int_\Lambda^1 us\, ds \approx \frac{\Lambda^2}{2} + \int_\Lambda^1 us\, ds. \tag{S16}$$

We now exploit the fact that $\langle u \rangle = 1/2$ in a symmetric integral about the midpoint of the transition zone [45]. Combining and simplifying gives the approximation for: $\rho(T)$

$$\rho(T) = R\sqrt{\frac{2\bar{u}}{1 + \Lambda^2}}. \tag{S17}$$

The infiltration is given by:

$$\Psi = \frac{R}{\rho_0}\sqrt{\frac{2\bar{u}}{1 + \Lambda^2}} - 1. \tag{S18}$$

Since we assume that the transition width is unknown, we can bound $\Psi$ by considering $\Lambda = 0, 1$.

#### Model calibration using spatial growth patterns

Coordinates of all spots within a well were calculated based on Supplementary Fig. 1A. We calculated what percentage of the total cells seeded per well ($2 \times 10^5$) fall within a spot. This was used to obtain the distribution of initial cell counts per spot (Supplementary Fig. 1B). To calculate the confluence of each spot at the time of seeding, we assumed an average cell diameter of 26.77 $\mu$m and a spot diameter of 350 $\mu$m (Supplementary Fig. 1C).

Cells detected within each spot three days post-seeding were assigned to bins within a grid $\tilde{G} \in R^{20,20}$, whereby each entry of $\tilde{G}$ is associated with its corresponding xy-coordinates $O \in R^{20,20,2}$. The number of cells within each bin was averaged across all replicates of a given ECM, $p$. Let $c := max_{p,i,j}(\tilde{G}_{i,j}(p))$ be the number of cells per bin at 100% confluence. For all $p$, $\tilde{G}(p)$ was divided by $c$ to obtain the $G(p)$ – the spatial distribution of confluence for $p$. For all entries of $G$, we calculated the euclidean distance, $D_{i,j}$, of its coordinates $O_{i,j}$ to the origin $(0, 0)$ of the MEMA spot. For all bins $B_t := \{i, j | t - e \leq D_{i,j} \leq t + e\}$ within distance $t$ to the origin, we then calculated the average confluence associated with that distance as:

$$\frac{1}{|B_t|} * \sum_{i,j \in B_t} G_{i,i} \tag{S19}$$

This was then compared to the confluence obtained from the simulations (Table 1) using the Wasserstein metric. 2.29% simulations fell within the bottom 0.05% quantile of Wasserstein distances and were considered representative for cellular growth on at least one ECM.

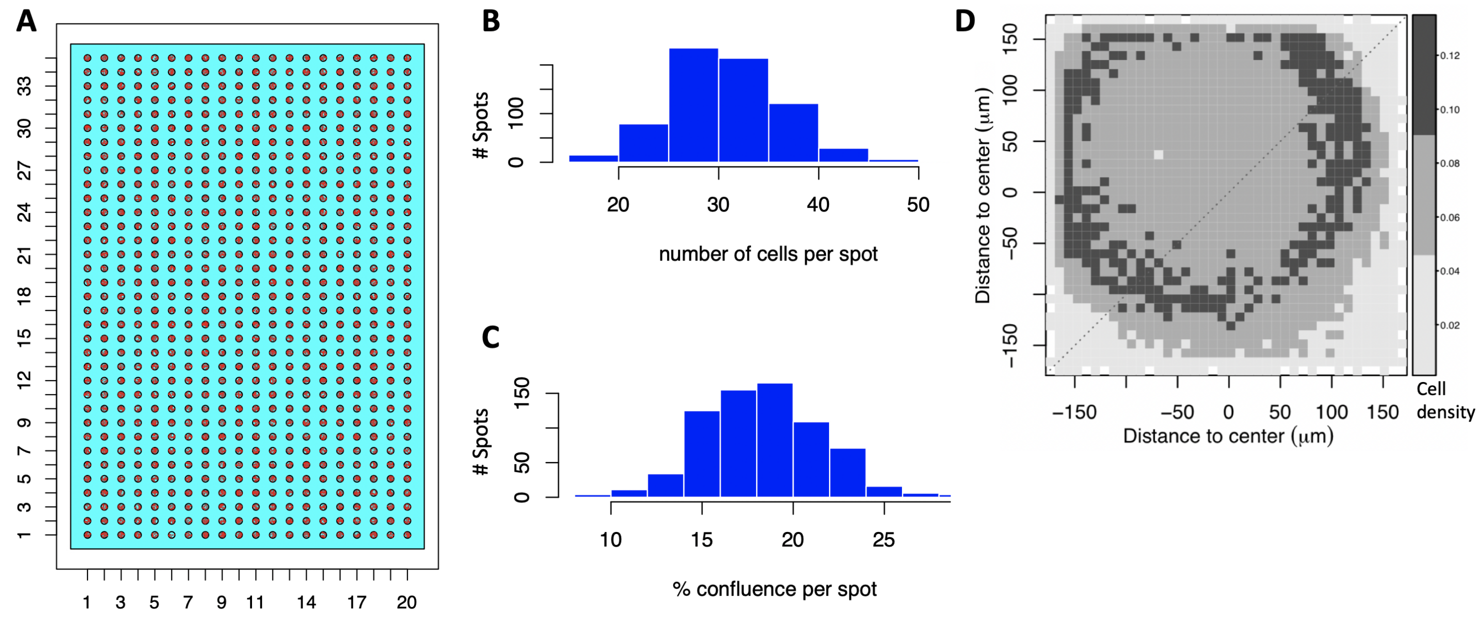


**Supplementary Figure 1. MEMA array platform design**. (A-C) Parameters inferred at the beginning of the MEMA assay. (A) Number of spots per row and column of a single well: $35 \times 20$; Diameter of a single spot: 350 $\mu$m; Horizontal and vertical distances between centers of two adjacent spots: 900 $\mu$m. Each spot image has 1600 x 1600 pixels, whereby 1 pixel = 321 nm. (B) Expected number of cells overlapping with a spot was calculated from parameters in (A), assuming $2 \times 10^5$ cells were distributed uniformly throughout the entire well [11, 60]. (C) Expected confluence per spot calculated from (B), assuming an average cell diameter of 26.77 $\mu$m. (D) Density distribution of spatial growth patterns is shown in aggregate across all spots in (A). Cell densities were measured at the end of the MEMA assay, after exposing HCC1954 cells to HGF for three days.

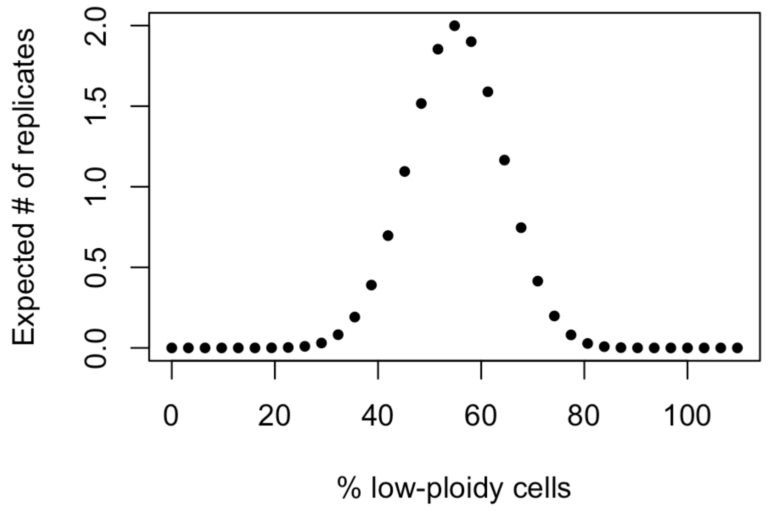


**Supplementary Figure 2. Hypothetical example of the influence of population heterogeneity on reproducibility across MEP replicates**. Probability of sampling given % low-ploidy cells when drawing 31 cells from a polyploid population (x-axis). Maximum probability shows true hypothetical incidence of low-ploidy cells (55% of the population). Probability was multiplied by the number of replicates (14) to obtain the values along the y-axis.

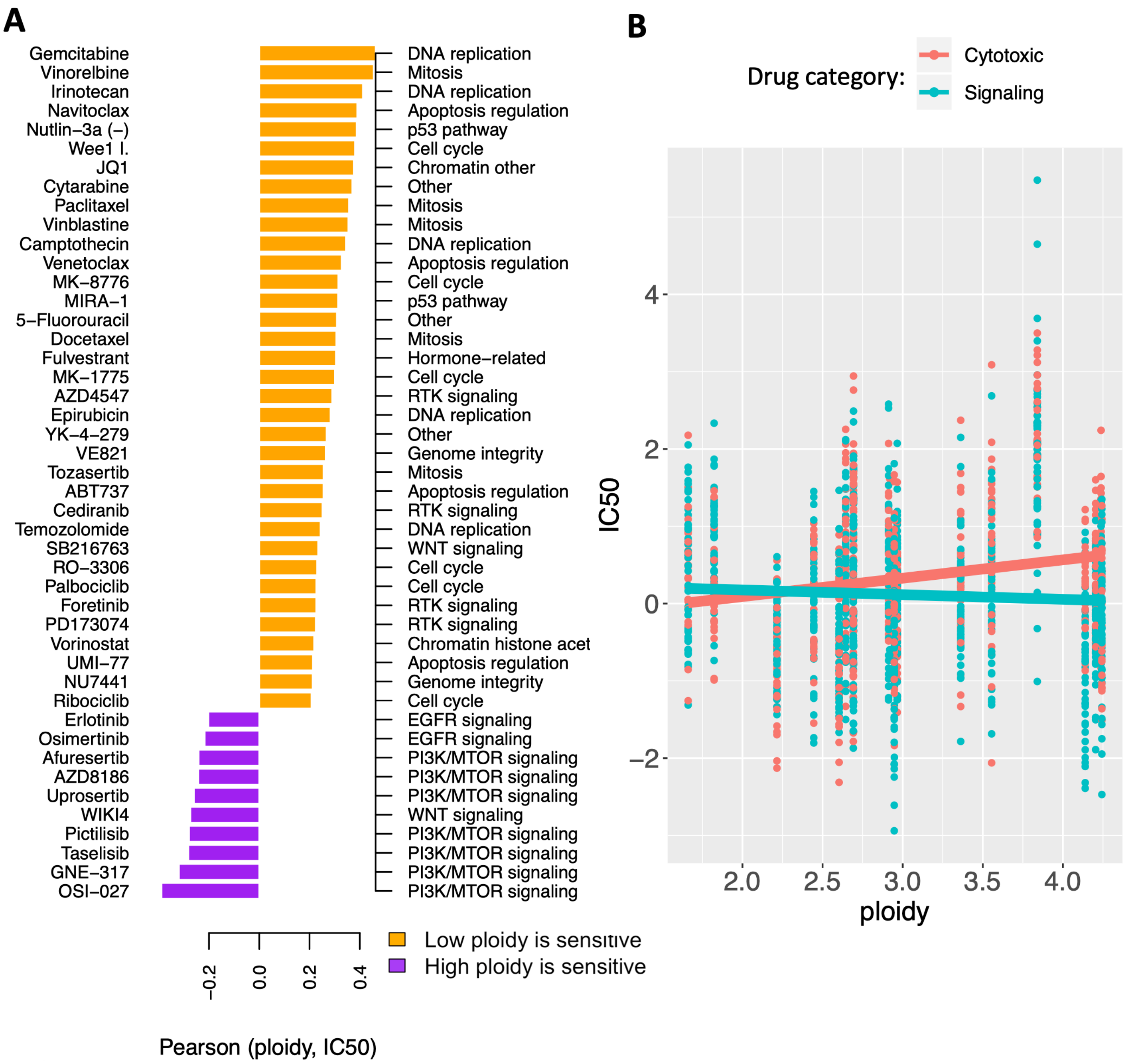


**Supplementary Figure 3. Validation of ploidy as biomarker of drug sensitivity across breast cancer cell lines in an independent dataset**. (A) High-ploidy breast cancer cell lines are resistant to cytotoxic drugs, but tend to be more sensitive to inhibitors of mTOR, EGFR and WNT signaling pathways. In a multivariate regression model of drug sensitivities molecular subtype alone (Fig. 1B), could explain 11% of the variability in IC50 values across cell lines (adjusted R-square = 0.109; $p < 1e\text{-}5$). Including ploidy into the model did minimally improve its predictive accuracy (adjusted R-square = 0.116; $p < 1e\text{-}5$). (B) However, an interaction term between ploidy and drug category increased accuracy to explain 14% of variability in drug sensitivity across cell lines (adjusted R-square = 0.142; $p < 1e\text{-}5$).

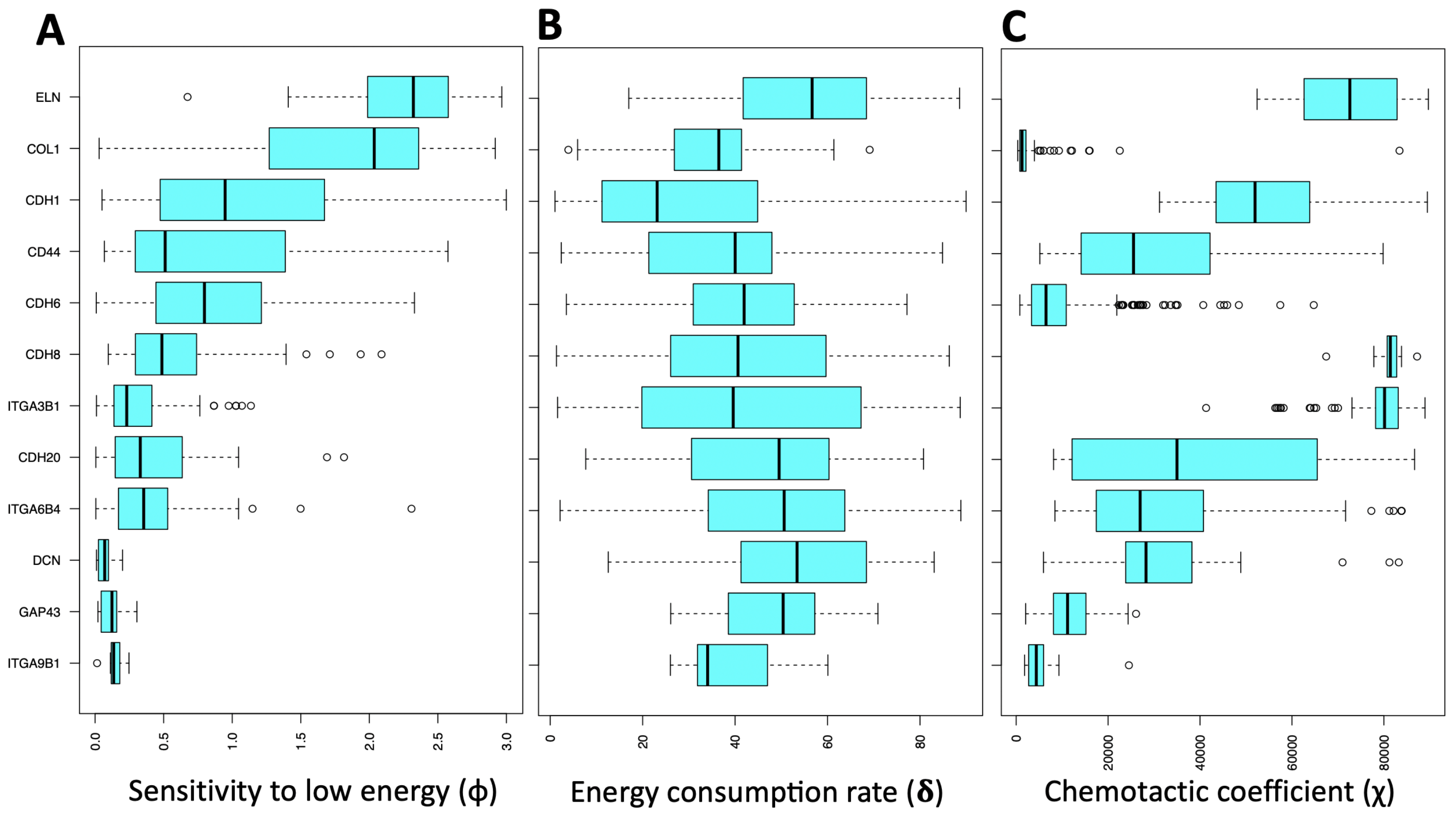


**Supplementary Figure 4. Posterior parameter distributions per ECM**. Best parameter fits for sensitivity to low energy (A), energy consumption rates (B) and chemotactic/haptotactic coefficients (C), stratified by the ECM they best explain.

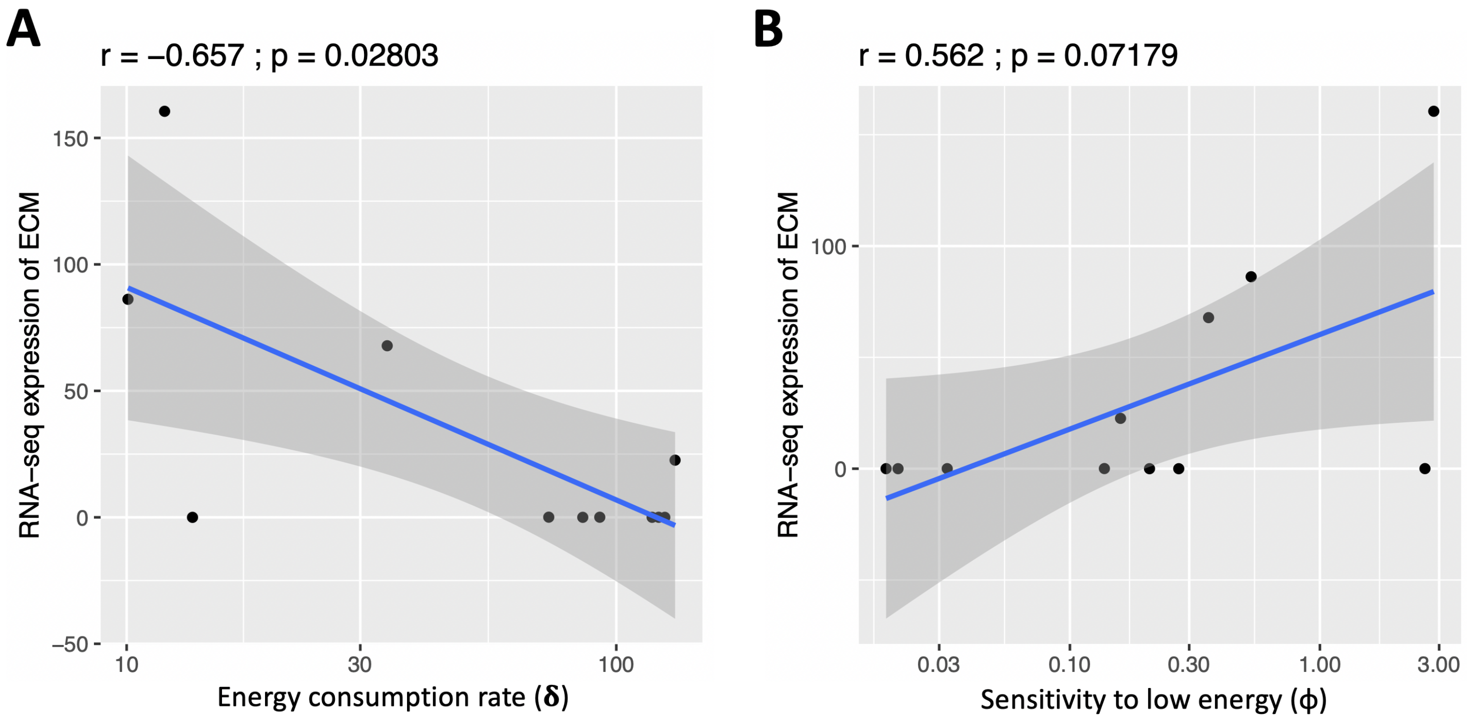


**Supplementary Figure 5. Correlation between inferred model parameters and RNA-seq derived signatures**. Model parameters $a$ (energy consumption rate; A) and $\phi$ (sensitivity to low energy; B) vary across ECMs (x-axis), and this variability correlates with the expression of the corresponding ECM (y-axis). RNA-seq derived signatures were not used in any way to infer any model parameter.

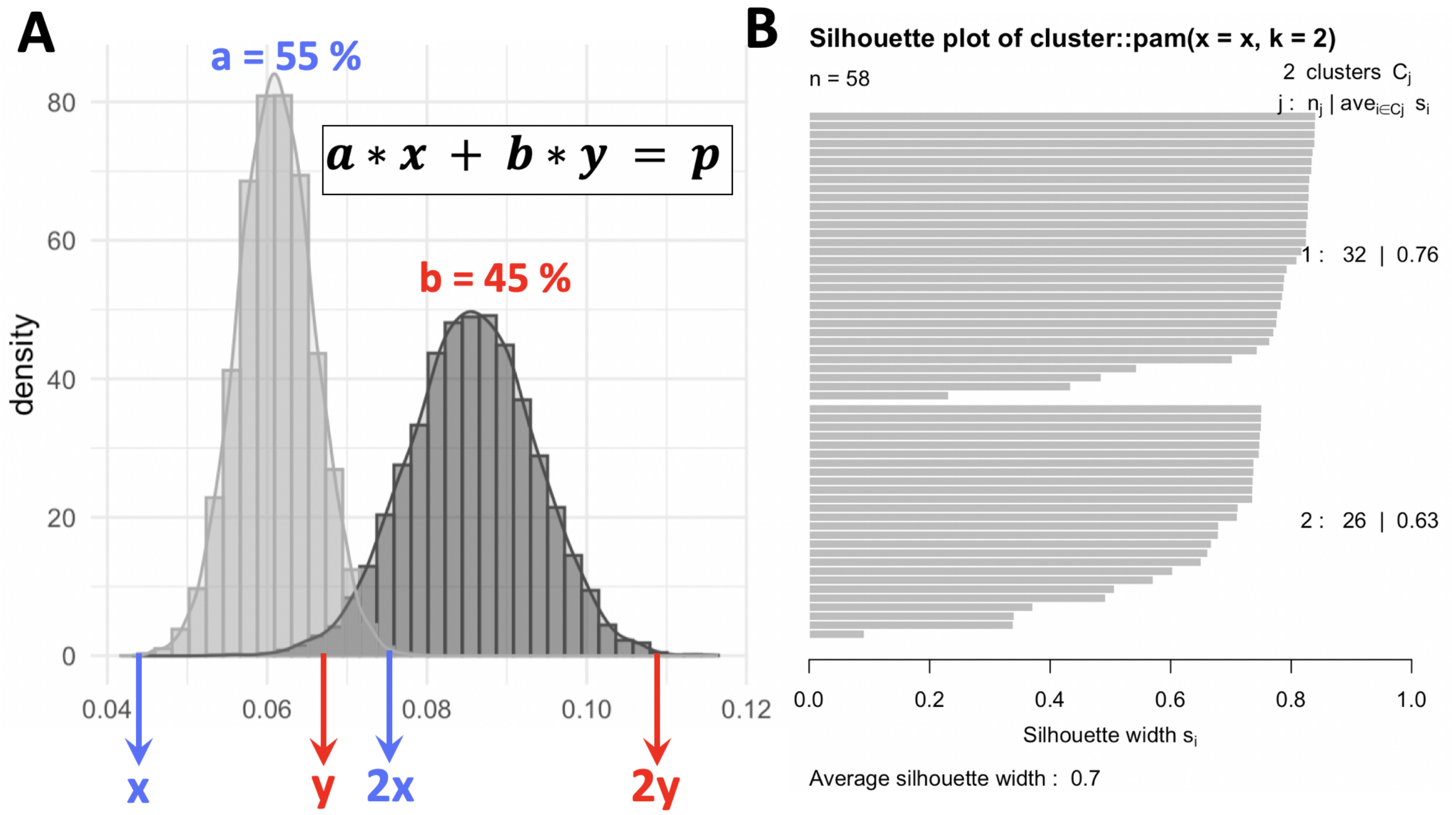


**Supplementary Figure 6. Estimating the ploidies of co-existing clones in the HCC1954 cell line**. (A) The bimodal distribution of DAPI content (x-axis) across EdU+ HCC1954 cells (y-axis) was used to infer the DNA contents of two clones thought to co-exist in the cell line (here denoted x and y respectively). The distribution informs the location of y as a function of x and the relative proportions of the two populations can also be read from the bimodal distributions of their S-phase cells. $P$ denotes the population-average ploidy of the HCC1954 cell line (4.2). Solving for $x$ yields ploidies 3.17 and 5.47 for the low and high-ploidy clone respectively. (B) Silhouette coefficient of each cell's assignment to one of the two clusters in (A). 18.9% of assignments (11 out of 58 cells) are ambiguous (silhouette coefficient < 0.6).

| Parameter | Pearson coefficient | p-value |
|---|---|---|
| Metabolism of vitamins and cofactors | 0.661 | 0.001 |
| Metabolism of water-soluble vitamins and cofactors | 0.597 | 0.005 |
| Cytochrome P450 - arranged by substrate type | 0.564 | 0.010 |
| Hyaluronan metabolism | 0.560 | 0.010 |
| Hyaluronan uptake and degradation | 0.560 | 0.010 |
| Glycerophospholipid biosynthesis | 0.556 | 0.011 |
| O-linked glycosylation of mucins | 0.553 | 0.011 |
| Synthesis of PC | 0.546 | 0.013 |
| Termination of O-glycan biosynthesis | 0.529 | 0.017 |
| Abacavir transport and metabolism | 0.520 | 0.019 |
| Advanced glycosylation endproduct receptor signaling | 0.513 | 0.021 |
| Purine catabolism | 0.513 | 0.021 |
| EPH-ephrin mediated repulsion of cells | 0.510 | 0.022 |
| The canonical retinoid cycle in rods (twilight vision) | 0.506 | 0.023 |
| Keratinization | 0.505 | 0.023 |
| RA biosynthesis pathway | 0.501 | 0.024 |
| Dectin-2 family | 0.501 | 0.025 |
| POU5F1 (OCT4), SOX2, NANOG activate genes related to proliferation | 0.490 | 0.028 |
| Biological oxidations | 0.486 | 0.030 |
| GPVI-mediated activation cascade | 0.472 | 0.036 |
| Attenuation phase | 0.470 | 0.037 |
| Lipoprotein metabolism | 0.468 | 0.037 |
| CD28 dependent PI3K/Akt signaling | 0.466 | 0.038 |
| Activation of SMO | -0.464 | 0.040 |
| Protein-protein interactions at synapses | -0.446 | 0.049 |
| Ephrin signaling | 0.445 | 0.049 |
| CD28 co-stimulation | 0.444 | 0.050 |

**Supplementary Table 1.** REACTOME pathways correlated to ploidy across 20 breast cancer cell lines from CCLE.

| Name | Pathway | Drug Category |
|---|---|---|
| Doxorubicin | Cytotoxic | Cytotoxic |
| Gemcitabine | Cytotoxic | Cytotoxic |
| Paclitaxel | Cytotoxic | Cytotoxic |
| Irinotecan | DNA replication | Cytotoxic |
| Ixabepilone | Microtubuli | Cytotoxic |
| Docetaxel | Cytotoxic | Cytotoxic |
| Ms275 | Cytotoxic | Cytotoxic |
| CPT-11 | Cytotoxic | Cytotoxic |
| Taxol | Mitosis | Cytotoxic |
| Topotecan | DNA replication | Cytotoxic |
| (Z)-4-Hydroxytamoxifen | Cytotoxic | Cytotoxic |
| Trichostatin A | HDAC Inhibitor | Cytotoxic |
| Vorinostat | Cytotoxic | Cytotoxic |
| Geldanamycin | Cytotoxic | Cytotoxic |
| Oxaliplatin | DNA replication | Cytotoxic |
| Vinorelbine | Mitosis | Cytotoxic |
| Cisplatin | DNA replication | Cytotoxic |
| Pemetrexed | Replication | Cytotoxic |
| Bortezomib | Protein stability and degradation | Cytotoxic |
| Carboplatin | DNA | Cytotoxic |
| 17-AAG | Cytotoxic | Cytotoxic |
| Nutlin 3a | MDM2 inhibitor, Apoptosis | Cytotoxic |
| CGC-11047 | DNA replication | Cytotoxic |
| 5-FU | Other | Cytotoxic |
| Imatinib | Apoptosis | Cytotoxic |
| Crizotinib | ALK/HGFR Inhibitor | Signaling |
| PS-1145 | IKB kinase | Signaling |
| Rapamycin | Signaling | Signaling |
| Omipalisib | Signaling | Signaling |
| AG1478 | Signaling | Signaling |
| PD184352 | MEK1/2 inhibitor | Signaling |
| Sirolimus | PI3K/MTOR signaling | Signaling |
| Cetuximab | EGFR | Signaling |
| Gefitinib | EGFR signaling | Signaling |
| Pd0325901 | MEK Inhibitor | Signaling |
| Trametinib | ERK MAPK signaling | Signaling |
| AG1024 | MAPK/ERK2 signaling | Signaling |
| Lapatinib | HER1/EGFR/ERBB1 Inhibitor | Signaling |
| Erlotinib | EGFR signaling | Signaling |

**Supplementary Table 2.** Drug classification.

| **Parameter** | ***Dimensional symbol*** | **Value** | **Units** | ***Dimensionless symbol*** | **Range** | **Identity** |
|---|---|---|---|---|---|---|
| Maximal growth rate | $\boldsymbol{\lambda}$ | 0.564 | $day^{-1}$ | | | |
| Radius of the dish | **Ř** | 1750 | µm | R | [4, 93] | |
| Energy consumption rate | $\boldsymbol{\delta}$ | 57 ⸸ | $cells^{-1}\ day^{-1}$ | a | [2, 160] | $\boldsymbol{\delta} = \lambda * a$ |
| Energy diffusion | $\boldsymbol{\Gamma_E}$ | 9500 ⸸ | $\mu m^2\ day^{-1}$ | $\eta$ | [0.01, 968] | $\Gamma_E = \chi * \eta$ |
| Simulation time | **T** | 3 | days | $\boldsymbol{\tau}$ | 1.69 | $T = \boldsymbol{\tau} / \lambda$ |
| Chemotactic coefficient | $\boldsymbol{\chi}$ | 5500 ⸸ | $\mu m^2\ day^{-1}$ | | | $\chi = \lambda * L^2$ |
| Characteristic length | | | | L | [19, 399] | L = Ř / R |

**Supplementary Table 3.** Conversion between dimensional and non-dimensional model parameters. Prior distributions of dimensionless parameters were drawn from the indicated ranges while ensuring uniform priors of dimensional parameters. The spatial growth patterns resulting from these prior distributions were compared to the MEMA data to obtain the maximum posterior estimates for dimensional parameters shown here.

8. Pienta KJ, Hammarlund EU, Axelrod R, Brown JS, Amend SR. Poly-aneuploid cancer cells promote evolvability, generating lethal cancer. Evolutionary Applications;n/a(n/a). _eprint: https://onlinelibrary.wiley.com/doi/pdf/10.1111/eva.12929. Available from: https://onlinelibrary.wiley.com/doi/abs/10.1111/eva.12929.

14. Growth of human diploid cells (strain MRC-5) in defined medium; replacement of serum by a fraction of serum ultrafiltrate. - PubMed - NCBI;. Available from: https://www.ncbi.nlm.nih.gov/pubmed/422676.

41. Sobol IM. Global Sensitivity Indices for Nonlinear Mathematical Models; 2004. .

60. Lin CH, Lee JK, LaBarge MA. Fabrication and use of microenvironment microarrays (MEArrays). Journal of Visualized Experiments: JoVE. 2012 Oct;(68).
